# Artoo-Detoo: What imitating a Star Wars droid reveals on allospecific vocal imitation in parrots and starlings

**DOI:** 10.1101/2024.10.10.617563

**Authors:** Nick C.P. Dam, Henkjan Honing, Michelle J. Spierings

## Abstract

Vocal production learning is a widespread and remarkable phenomenon. However, the underlying mechanisms of allospecific vocal imitation are poorly understood. Parrots and corvids are well known for their imitation abilities, but imitation accuracy has not yet been studied across species in a comparative context. We compared imitation accuracy between nine parrot species and European starlings based on imitation of monophonic and multiphonic sounds, produced by the Star Wars droid R2-D2. Our results show that starlings were better at imitating multiphonic sounds than parrots. However, no difference in imitation accuracy was found between parrots and starlings when imitating monophonic sounds. The differences in imitating accuracy are likely based on a physiological difference in syrinx anatomy, rather than on perceptual or cognitive differences between starlings and parrots. Between our nine tested parrot species, we found that parrots with larger brains (e.g., African greys and amazon parrots) were less accurate at imitating monophonic sounds than smaller brained budgerigars and cockatiels. The use of citizen science in our study shows the great potential this has to expand data collection beyond traditional studies. These findings contribute to a deeper understanding of the evolution of communication complexity in vocal learners.

**Significance Statement:** Vocal imitation is a hallmark of complex communication in animals, yet the mechanisms driving species differences remain poorly understood. By comparing vocal imitation across nine parrot species and European starlings, we demonstrate, for the first time, that differences in syrinx anatomy, rather than cognitive or perceptual capacities, underlie species-specific variation in imitation accuracy. Contrary to traditional expectations, smaller-brained parrots outperformed larger-brained species in monophonic sound imitation. These findings challenge assumptions that brain size is directly linked to imitation accuracy and offer insights into the evolution of complex communication. We used citizen science to create a uniquely large dataset of species imitating the same source, demonstrating the powerful potential of this approach to broaden research in comparative cognition and vocal learning.

## Introduction

Vocal production learning, a complex and intriguing phenomenon, has independently evolved across various taxa, including elephants, bats, pinnipeds and cetaceans (1). However, vocal learning is particularly apparent in avian species varying from songbirds (2) to parrots, hummingbirds and some other bird species (3, 4). While songbird vocal learning has been well-characterized through iconic models like the zebra finch (*Taeniopygia guttata*) (5) and white-crowned sparrow (*Zonotrichia leucophrys*) (6), comparative studies exploring vocal learning processes in parrots remain scarce (3). Moreover, direct comparisons of vocal production learning between parrots and songbirds are currently lacking with regards to vocal production complexity (7). The complexity of vocal production learning is often discussed in relation to repertoire size, but less so in terms of the structure of the produced sounds. Nevertheless, certain birds may imitate a wide variety of sounds including human speech (8-10), environmental sounds (11) but also artificial sounds like chainsaws (12).

Imitating sounds from different species or anthropogenic sources, or allospecific vocal imitation, allows birds to add more vocal complexity to their vocal repertoire (13). This enhanced vocal variation can function to deter predators by reproducing allospecific alarm calls when disturbed at the nest, as in spotted bowerbirds (*Ptilonorhynchus maculatus*) (14), to imitate the vocalizations of host species during brood parasitism (cuckoos) (15) and to create complex songs to attract mates as a form of sexual selection in European starlings (*Sturnus vulgaris*) (16). However, the quality of allospecific vocal imitation is usually compared between one species and the model sounds, but not across different species imitating the same model sound (17-20).

Specifically, a comparison between species from different clades and with different vocal systems could give insight into the role of cognitive abilities and/or physiological structures in vocal imitation accuracy. For example, songbirds (such as starlings) and parrots (like budgerigars; *Melopsittacus undulatus*) are both vocal learners with extensive repertoires, but also have distinct syrinx structures (SI Appendix, Fig. S1). Starlings have independent control over the tracheobronchial muscles at both sides of the syrinx which allow them to produce two sounds at the same time (21). In contrast, parrots do not have this independent control over the syrinx’ muscles and thus lack this biphonation ability (22-24). Moreover, songbirds and parrots also differ in the neurological structures underlying vocal production learning (25). Vocal learning is neurologically controlled by a well described “song system” in songbirds. In parrots, similar neurological structures have been found in an area called the “core” region. However, the nuclei surrounding the core region in the so called “shell” region, which is unique to parrots, have been suggested to play an important role in vocal imitation. Variation in relative size between these core and shell regions has been hypothesized to be related to variation in the complexity of vocal production learning between parrot species (25).

This study presents a dataset of ten different species all imitating the exact same, complex, sound model: R2-D2 the renowned droid from Star Wars. We analyzed allospecific vocal imitation accuracy in eight different vocal units. R2-D2’s sounds were created by Ben Burtt using an ARP 2600 modular synthesizer in combination with Ben Burtt’s own voice (26). The vocabulary of R2-D2 can be distinguished in two types of sounds: monophonic and multiphonic sounds. Whilst monophonic sounds are relatively simple synthesized bleeps and clicks, the multiphonic sounds were generated using the ring modulator of the ARP 2600. A ring modulator is a signal processing device that takes two input signals—a carrier and a modulator—and multiplies them to produce a complex output. This multiplication creates new frequencies known as “sidebands,” which are the sum and difference of the frequencies of the input signals. For example, if the carrier signal has a frequency of 1,500 Hz and the modulator signal is 400 Hz, the output would consist of two sine waves: one at 1,900 Hz (sum of the two frequencies) and one at 1,100 Hz (difference). These sidebands reflect the combined spectral information from the inputs. If the modulator signal is more complex, the output produces mirrored spectral patterns (i.e. *spectral contours*; ten Cate & Honing (27): Fig. 29.2) around the carrier frequency. See SI Appendix, Fig. S2*F* for an example of such a mirrored multiphonic sound.

Nine parrot species and European starlings were analyzed to answer three key questions 1) How do parrots and starlings differ in their imitation capabilities? 2) How do different parrot species compare in their ability to replicate these complex sounds? 3) What are the differences in performance between multiphonic and monophonic sounds? Our study gives insights in a broader vocal imitation framework. Understanding the nuances of allospecific imitation in birds provides valuable insights into the cognitive and neural mechanisms driving vocal learning. By comparing imitation accuracy between two distinct vocal systems—the syrinx of parrots and starlings— and species with different cognitive abilities, we highlight key differences in adaptability and learning diversity across species.

## Results

### Starlings and parrots imitate monophonic and multiphonic units differently

We used citizen science to collect 107 videos of nine different parrot species and eight videos of European starlings (115 videos in total) imitating R2-D2 sounds uploaded by parrot owners on various social media platforms (YouTube, TikTok and Instagram). We annotated each sound on an elemental (single sounds) and unit level (combination of multiple elements; SI Appendix, Fig. S2) in Praat (28). A total of 2130 imitations of 98 different units were imitated by the birds. We selected the most frequently imitated monophonic units (units: “H”, “M”, “N” and “P”; SI Appendix, Fig. S2 *A*–*D*) and multiphonic units (units: “X”, “Y”, “C” and “W”; SI Appendix, Fig. S2 *E*–*H*). This resulted in a comparison of 419 monophonic and 463 multiphonic units imitated by birds (n = 101) to the original model R2-D2 units.

Spectrograms show that starlings were able to imitate multiphonic sounds and reproduce the two mirrored spectral contours created by ring modulation (Fig. 1 *D* and *E*). Six out of eight starlings imitated these mirrored contours when imitating multiphonic sounds. On the other hand, parrots always copied only the fundamental frequency of multiphonic units and never imitated the mirrored contours (Fig. 1 *D* and *F*).

**Figure 1.**
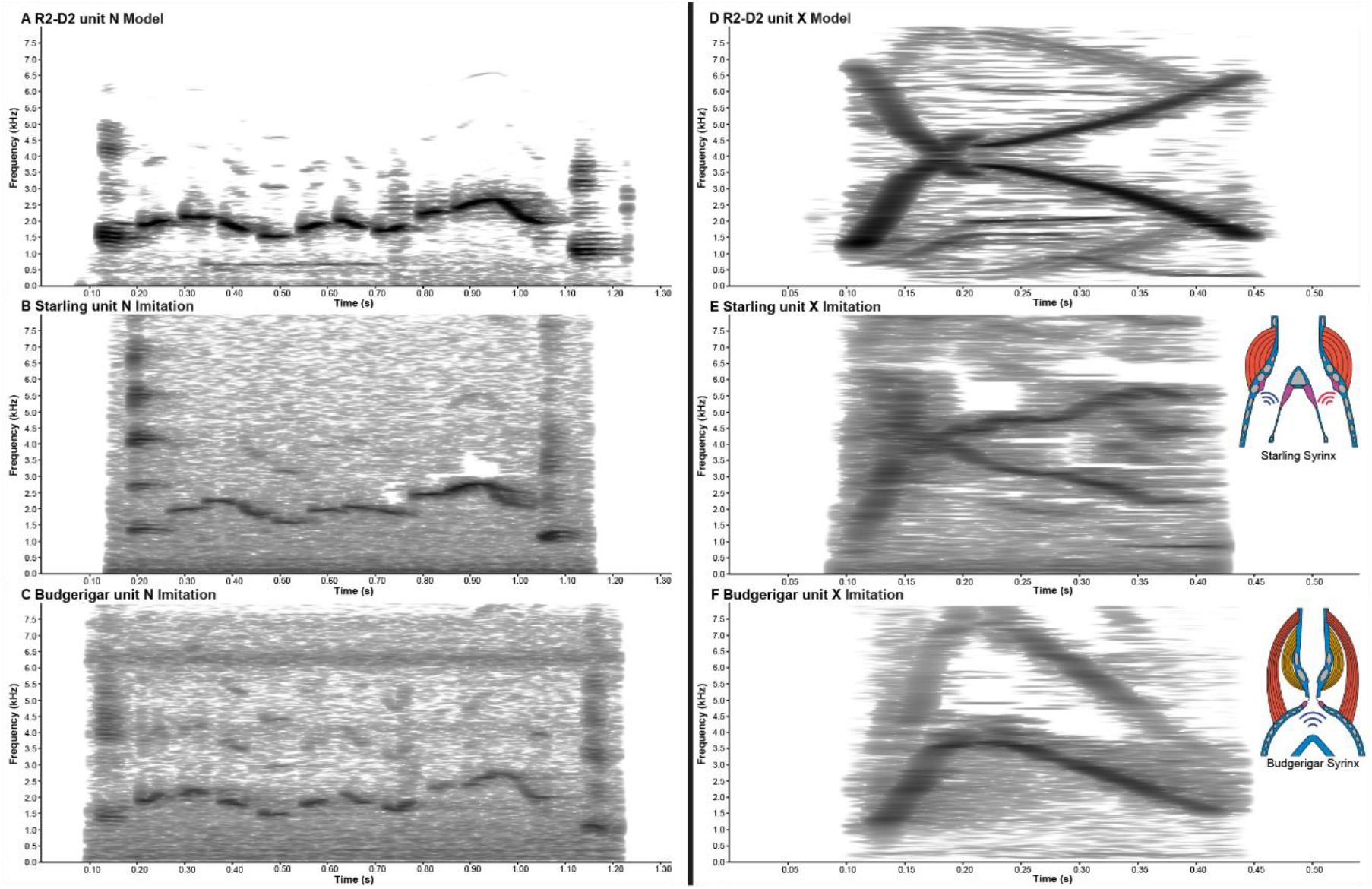
Spectrograms comparing the model R2-D2 sounds to imitations of starlings and parrots. (*A*) Example of a monophonic R2-D2 unit: “N”. (*B*) Imitation of unit “N” by a starling. (*C*) Imitation of unit “N” by a budgerigar. (*D*) Example of a multiphonic R2-D2 unit: “X”. (*E*) Imitation of unit “X” by a starling. (*F*) Imitation of unit “X” by a budgerigar. Syrinxes show the differences in the sound production mechanism with starlings being able to produce two sounds at the same time and parrots only one sound (see SI Appendix, Movie S2 for sound examples).

### Starlings imitate multiphonic units more accurately

To quantify this difference, we compared imitation accuracy between species by calculating dissimilarity scores for each unit using the dynamic time warping algorithm of Luscinia (29). Dissimilarity scores were calculated based on weightings of several spectro-temporal parameters (SI Appendix, Table S1 and methods for more details). A lower dissimilarity score indicates a more accurate imitation of the R2-D2 sound unit.

Imitation accuracy between species varied both between and within monophonic and multiphonic units (Fig. 2). Multiphonic units were in general more difficult to imitate accurately (mean: 8.758; range: 1.338 – 13.581) than monophonic ones (mean: 0.992; range: 0.103 – 7.791). Variation was also found in how accurately each unit was imitated within both types of R2-D2 units. For example, unit “P” was more accurately imitated than unit “H”, and unit “C” was more difficult to imitate than unit “W”, within the monophonic and multiphonic categories, respectively (Fig. 2 and SI Appendix, Fig. S2). Dissimilarity scores also varied within species on an individual level (SI Appendix, Fig. S3).

**Figure 2.**
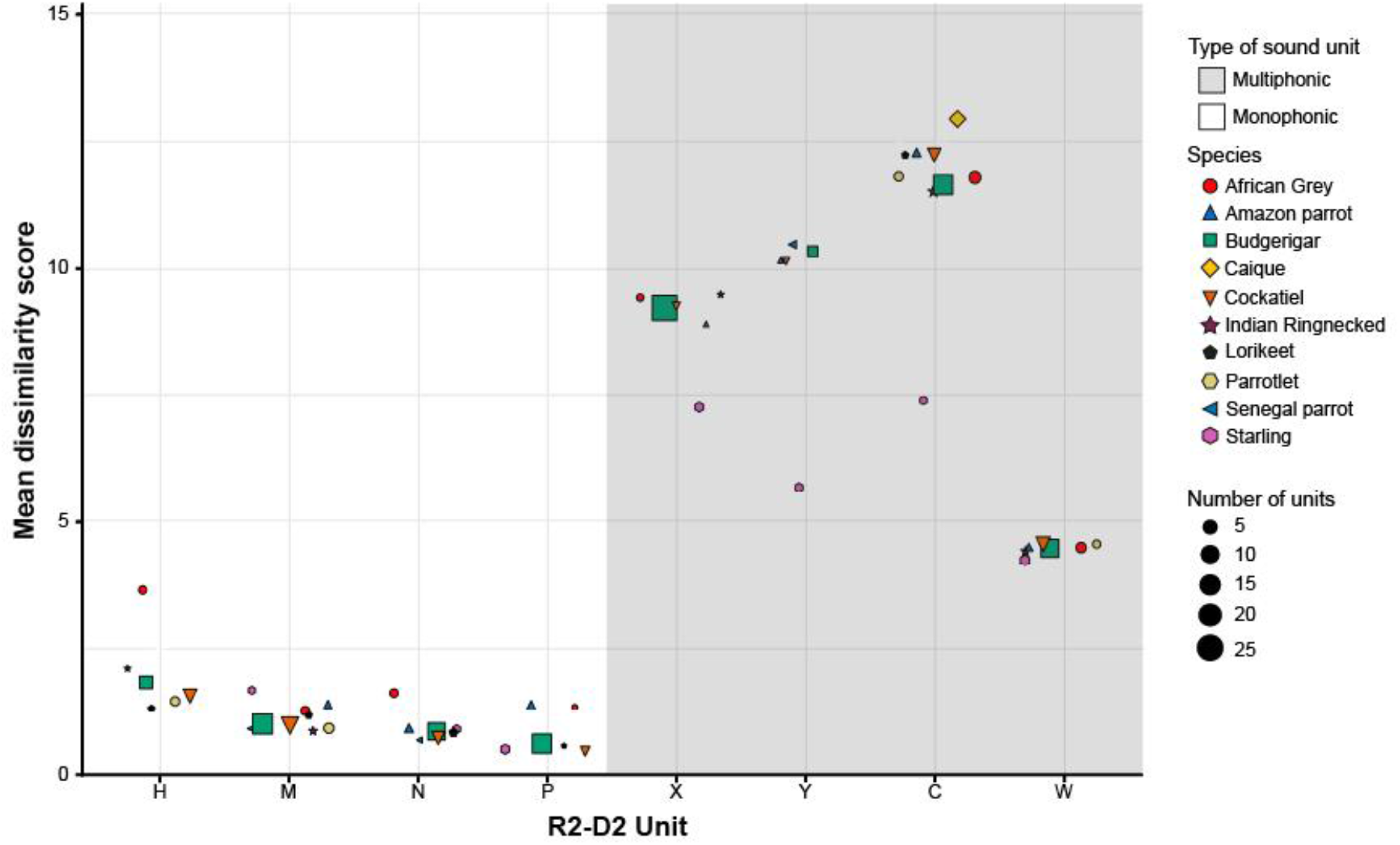
Imitation accuracy of monophonic and multiphonic R2-D2 units per species. Mean dissimilarity scores per unit for each species are shown. Dissimilarity scores varied between each unit. Colors and shapes indicate the different species. Scores in the white and grey background correspond to monophonic and multiphonic units, respectively. Sizes of dots are related to the number of imitated units per species.

To test whether starlings imitated multiphonic sounds more accurately than parrots we created a Linear Mixed Model (LMM). We found that dissimilarity scores were explained by an interaction between group (starlings vs. parrots) and monophonic or multiphonic units (LMM: -3.240 ± 0.293, *t* = -11.042, *p* < 0.001; SI Appendix, Table S2). A Post-Hoc test revealed that starlings imitated multiphonic R2-D2 units more accurately than parrots (Tukey: 2.771 ± 0.284, *t* = 9.759, *p* < 0.001; Fig. 3, SI Appendix, Fig. S4 and Table S3). Furthermore, multiphonic units were imitated less accurately than monophonic units by both parrots (Tukey: 2.630 ± 0.284, *t* = 9.271, *p* < 0.001; Fig. 3) and starlings (Tukey: 3.099 ± 0.286, *t* = 10.820, *p* < 0.001; Fig. 3) confirming that multiphonic units are more difficult to imitate than monophonic ones. Parrots and starlings did not differ in imitation accuracy for monophonic R2-D2 units (Tukey: -0.469 ± 0.328, *t* = -1.431, *p* = 0.482; Fig. 3 and SI Appendix, Table S3).

**Figure 3.**
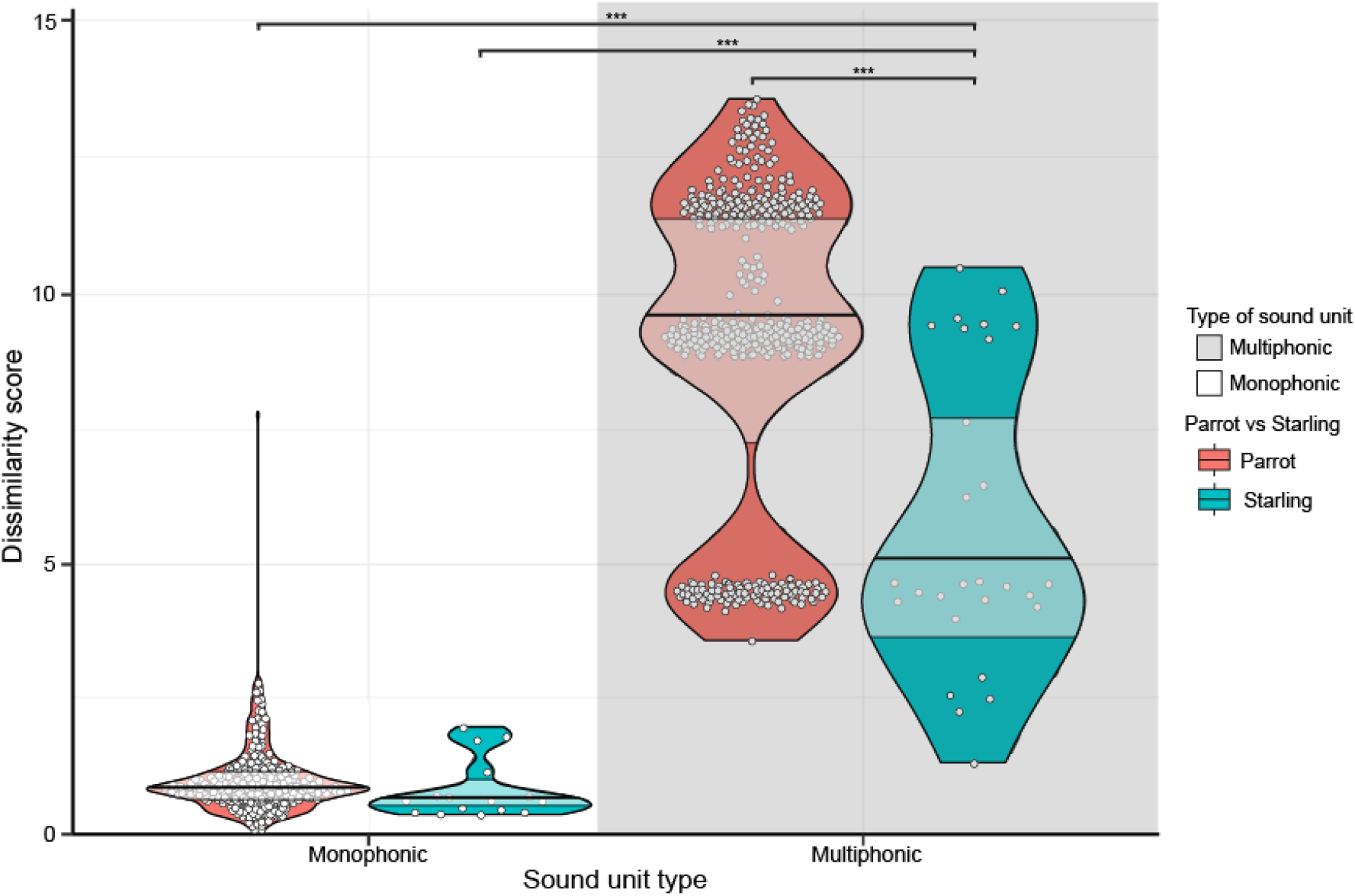
Starlings imitate multiphonic units more accurately than parrots, no difference in monophonic units. Starlings have lower dissimilarity scores for multiphonic units than parrots. Violin plots showing the distribution of the data. Horizontal lines and boxes correspond to medians and interquartile ranges. Each dot represents an individual imitating a sound unit. Dissimilarity scores were combined for all monophonic and all multiphonic units separately. Colors correspond to dissimilarity scores of starlings or parrots. Stars indicate significance levels (*** P < 0.001).

### Parrot species vary in imitation accuracy

Interestingly, the larger brained parrots, African greys (*Psittacus erithacus*, mean: 1.871; range: 0.872 – 7.791) and Amazon parrots (*Amazona* sp., mean: 1.198; range: 0.621 – 2.038), imitated the monophonic units the least accurate (SI Appendix, Fig. S4). In contrast, smaller brained species like budgerigars (mean: 0.909; range: 0.103 – 2.797), cockatiels (*Nymphicus hollandicus*, mean: 1.029; range: 0.267 – 2.769) and also starlings (mean: 0.839; range: 0.383 – 1.992) imitated monophonic units the most accurately. Moreover, budgerigars (Tukey: 1.400 ± 0.329, *t* = 4.258, *p* = 0.004; SI Appendix, Fig. S4) and cockatiels (Tukey: 1.613 ± 0.353, *t* = 4.573, *p* = 0.001; SI Appendix, Fig. S4) imitated monophonic units significantly more accurate than African greys, and cockatiels imitated more accurately than Amazon parrots (Tukey: 1.232 ± 0.340, *t* = 3.623, *p* = 0.041; SI Appendix, Fig. S4). Imitation of monophonic units by all other parrot species varied between the scores of African greys and budgerigars. Multiphonic units were imitated the least accurate by rainbow lorikeets (*Trichoglossus moluccanus*, mean: 12.233; range: 11.444 – 13.022) and caiques (*Pionites* sp., mean: 12.931; range: 12.332 – 13.581). Budgerigars (mean: 8.845; range: 3.590 – 12.868) and cockatiels (mean: 8.109; range: 4.323 – 13.474) had the lowest dissimilarity scores for multiphonic units amongst the parrots, similarly to the monophonic results (Fig. 2 and SI Appendix, S4). All other parrot species’ imitation accuracy varied in between the scores of budgerigars & cockatiels and lorikeets & caiques. In summary, parrot species varied in how accurately they imitated R2-D2 sounds with smaller brained parrots imitating R2-D2 sound more precisely than larger brained ones, and starlings imitated multiphonic sounds more accurately than parrots.

### Dendrograms confirm similarities between bird species and R2-D2

Finally, to corroborate our results we created dendrograms to hierarchically cluster species based on similarities between imitations (Fig. 4). As expected, starlings clustered together with the R2-D2 model in a dendrogram based on all multiphonic units, thus validating our previous results. In contrast, when comparing monophonic units, cockatiels’, budgerigars’ and parrotlets’ imitations were more similar to R2-D2, whereas starlings split off at the more basal end of the dendrogram and are thus less similar to R2-D2 (Fig. 4). Furthermore, African greys clustered at the most basal end of the monophonic dendrogram with the largest distance away from the R2-D2 model and therefore corroborating our earlier result of African greys imitating monophonic R2-D2 units less accurately. No overlapping results in species clustering were found between the monophonic and multiphonic dendrograms.

**Figure 4.**
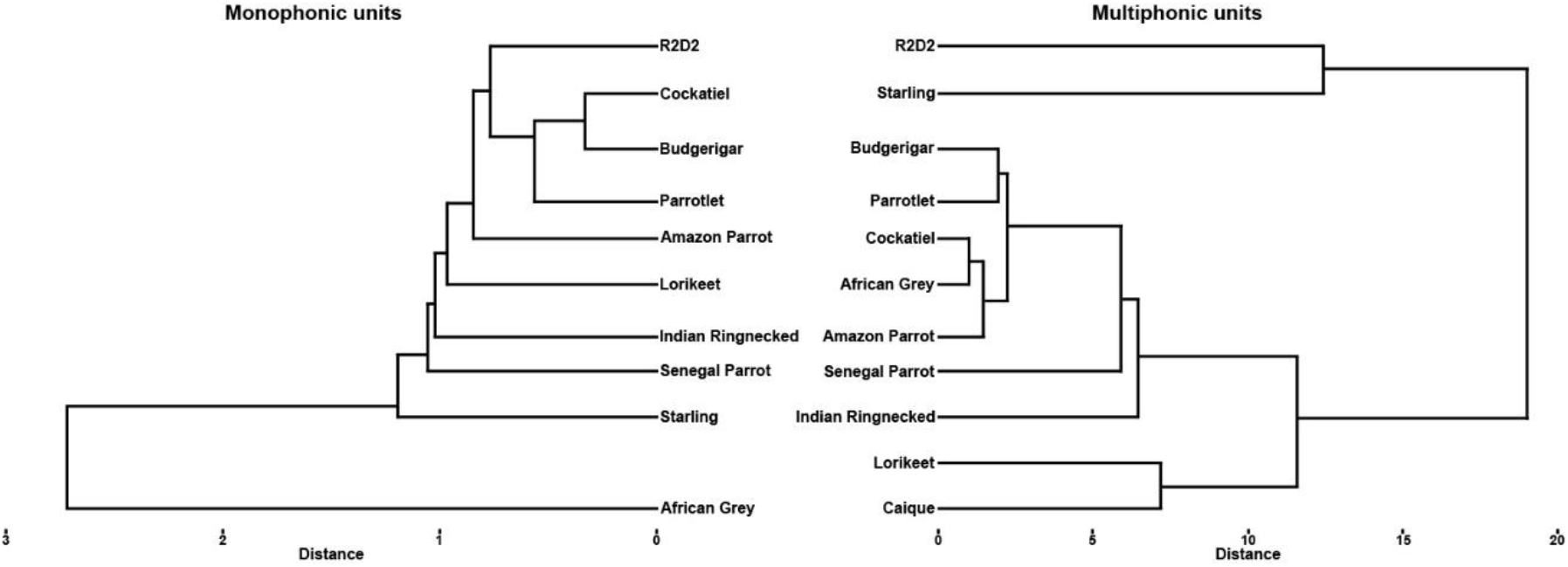
Dendrograms of imitation accuracy of R2-D2 units. Hierarchical representations of similarity of R2-D2 unit imitations between species. Left shows a dendrogram based on monophonic units while the right dendrogram is based on multiphonic units. Distances were calculated as Euclidean distances between all species and R2-D2.

## Discussion

The present study compared vocal imitation accuracy of monophonic and multiphonic sounds between nine parrot species and European starlings using R2-D2 sounds as the model. In general, monophonic units were better imitated than multiphonic units by all bird species. However, the starlings showed incredibly accurate imitation of multiphonic units, specifically compared to all parrot species. Some units were easier to copy than others, both within the monophonic and multiphonic sounds, which could be explained by some units potentially being more similar to the natural vocalizations of our tested species. Amongst the parrots, the smaller brained budgerigars and cockatiels were more accurate at imitating both monophonic and multiphonic sounds than the larger brained parrots, contradictory to results from studies on variation in core and shell nuclei size (25).

Starlings imitated monophonic sounds similarly to parrots, but were significantly more accurate at imitating multiphonic sounds than parrots. Interestingly, a physiological explanation seems most convincing due to the anatomical differences between starlings and parrots. The syrinx of starlings contains two vibrating sound sources called the lateral labia and medial labia (SI Appendix, Fig. S1). These labia can be independently controlled at both sides of the syrinx by the syringeal muscles thus making it possible for starlings to produce two pitches at the same time (22). In contrast, parrots lack the independent control of these labia at both sides of the syrinx. Therefore, parrots lack the biphonation ability that starlings have (22-24). Differences in perception or cognitive abilities can therefore be excluded as explanation for the observed differences in imitation accuracy since starlings did not differ in how accurate they imitated monophonic sounds from parrots. Although biphonation made it possible for starlings to imitate both the fundamental frequency and the mirrored spectral contours of the multiphonic units not all starlings imitated the two mirrored spectral contours. Two out of eight starlings did not copy the two spectral contours of any of the multiphonic R2-D2 sounds. We hypothesize that these two individuals did not receive sufficient training to accurately reproduce the fundamental frequency and the mirrored spectral contours. However, starlings learn environmental sounds that are frequent and similar to their own vocalizations and thus we could assume that the starlings heard the R2-D2 sounds frequently enough (14).

Parrot’s imitation accuracy of monophonic units showed an unexpected pattern. Chakraborty et al. hypothesized that parrots with relatively larger shell nuclei compared to core nuclei to have more complex vocal learning abilities. This relative size difference is larger in parrots with larger brains (like macaws, African greys and amazon parrots) which are also well known for their speech imitation abilities (8). Therefore, we expected these species to be more accurate at imitating monophonic sounds. However, in our study we found that parrots with larger brains, and also relatively large shell nuclei, imitated monophonic sounds significantly less accurate than budgerigars and cockatiels that have smaller shell regions and larger core regions. Parrots with smaller brains, however, have a smaller repertoire of imitated sounds (11) as expected by the results of Chakraborty et al. (25). It could therefore be possible that smaller brained parrots are more trained on imitating the few sounds they have learned and are thus more accurate than the larger brained parrots. However, budgerigars are also able to imitate speech at similar rates of parrots with larger shell regions (11) but do so by frequency modulating their ‘’warble’’ song and thus not producing new imitated sounds but modified natural vocalizations (10). Imitation accuracy could also not be related to the capacity of complex vocal learning and shell region size but instead the rate of learning could be more related to differences in shell size. Peach-fronted conures (*Eupsittula aurea*) for example have been shown to have more rapid vocal modification abilities compared to budgerigars which have smaller brains than conures (30). Furthermore, another possible explanation could be that the natural vocalizations of smaller brained parrots are more similar to R2-D2 units and thus easier to copy. However, vocal repertoires have not been extensively studied in all parrot species in the present study (7). Variation in imitation accuracy is also found between other bird species. Superb lyrebirds (*Menura novaehollandiae*) (17) and satin bowerbirds (*Ptilonorhynchus violaceus*) (18) are known to be incredible accurate imitators while the brown thornbill (*Acanthiza pusilla*) (19) and Icterine warbles (*Hippolais icterina*) (20) are less accurate at imitating sounds. Interestingly, the male bowerbirds imitate sounds to incorporate in their songs to attract females and the most accurate imitators are more attractive to female bowerbirds (18) whereas brown thornbills and Icterine warbles imitate alarm calls of other species (19, 20). Suggesting that vocal imitation accuracy varies depending on its function.

Multiphonic R2-D2 sounds were more difficult to imitate than monophonic sounds regardless of having biphonation abilities or not. Starlings imitated the mirrored spectral contours of the multiphonic sounds, however, it was still difficult to precisely imitate the sounds indicated by the higher dissimilarity scores. Moreover, large variation between imitation accuracy was found between different multiphonic units indicating that some units were easier to copy than others. Unit “W” consisted of only one rising mirrored formant and had the lowest dissimilarity scores. In contrast, unit “C” consisted of four rising and falling mirrored spectral contours and showed the least accurate dissimilarity scores of all sounds (SI Appendix, Fig. S2 *E*-*H*). Unit “X” and “Y” both had two rising and falling mirrored spectral contours and scores fell in between the previous two units suggesting that the amount of mirrored spectral contours is related to the complexity of the sounds. Similarly, variation in imitation accuracy was found between monophonic units but to a smaller degree than with the multiphonic units. Units with more rising and falling pitches (like unit “H”) were more difficult to imitate than units with a flatter pitch contour (like unit “P”; SI Appendix, Fig. S2 *A*-*D*).

One of the drawbacks of the present study it that the amount of training for each individual is unknown. Individual variation within each species was large, which could be an indication of different levels of auditory experience. However, most individuals within a species showed similar dissimilarity scores while a few had deviating (higher) scores than the rest suggesting that most individuals received similar amounts of training. Furthermore, it is also unknown how they were trained, i.e. only hearing playbacks of R2-D2 sounds versus also receiving positive rewards like food. We only have anecdotal stories of video descriptions mentioning that one of the parrots started to imitate R2-D2 sounds after listening to 3 hours of playbacks. In general, knowledge about the processes behind vocal learning of imitated sounds are largely lacking (31). Furthermore, usually only one species is compared with the respective model sound that is being imitated (17-20). However, based on our results and other studies (11) we can hypothesize that species with a wider vocal repertoire of imitated sounds may be less accurate at imitating sounds whereas species with smaller repertoires put more energy in producing more accurate sound copies rather than imitating more sounds. Moreover, male starlings may imitate sounds at higher accuracy so that females find their songs more attractive than male songs with less accurate imitations, similar to satin bowerbirds (18). The use of citizen science enabled us to compare imitation accuracy across ten different species with each other showing the potential citizen science has to increase the volume and diversity of data collection. Our study helped contributing to compare imitation accuracy across species however the difference in learning capacity between species is still unknown. Similarly to Schachner et al. (32), we collected videos from YouTube to conduct our study, however, citizen science could also be used to directly work together with companion parrot owners to conduct more detailed studies to have a deeper understanding behind the learning processes of allospecific imitation and the evolution of vocal complexity, like the initiatives as the bird singalong project (33) and many parrots project (34).

In conclusion, we showed that starlings are able to more accurately imitate multiphonic sounds than parrots due to differences in syrinx anatomy. Parrots and starlings did not differ in imitation accuracy of monophonic sounds. However, variation in imitation accuracy was found between parrot species which could potentially be related to differences in brain size. Furthermore, more studies are needed as to how parrots and starlings learn to imitate these complex sounds. What are the learning mechanisms behind vocal imitation and how do they differ from other forms of vocal production learning? Finally, other questions can be answered with the dataset we have in which we know the source in future studies: are differences in imitation accuracy related to difference in relative core and shell region size? Are there perceptual biases for certain acoustic dimensions that differ between species and could these be related to natural vocalizations? Which aspects of the model are copied (in an absolute fashion) and which are relatively imitated (e.g. transposed pitch but maintained pitch countour)? Are there temporal differences in imitating R2-D2 sounds between species? Answering these questions will contribute to our understanding of the evolution of complex communication, vocal learners, and getting a better understanding of the precursors of human language and music in animals (27).

## Materials and Methods

### Data collection

We collected videos of parrots and European starlings imitating R2-D2 sounds from YouTube, Instagram and TikTok. Search terms included “Parrot imitating R2-D2”, “Parrot R2-D2”, “Starling imitating R2-D2”, “Starling R2-D2” and the same search terms translated in other languages (Dutch, German, Spanish, Portuguese). Furthermore, search filter options like “most recent upload” were used to find more recent videos of better quality. Videos were downloaded using online video converters as MP4 files (YouTube: https://ssyoutube.com/en174YV/youtube-video-downloader; Instagram: https://snapinsta.app/nl; TikTok: https://snaptik.app/en1). Next, we converted each MP4 video to .wav format using an online file converter (https://cloudconvert.com/). We collected one video per individual since not all individuals had multiple videos available. For individuals that had more videos available online, we choose the video of highest quality if necessary (i.e. the other videos were of poor audio quality). We have no knowledge about how much training each bird had hearing the R2-D2 sound playbacks nor how they were trained, i.e. only playing back R2-D2 sounds or also rewarding them with food etc.

### Annotation

We identified and annotated each imitated R2-D2 sound in Praat (version 6.3.14) (28). Original R2-D2 sounds were first visually inspected on a spectrogram and annotated as letters for each individual sound unit. We hypothesized that most parrot and starling owners played back a video on YouTube to their bird titled “R2-D2 Sounds for bird mimicking” (https://www.youtube.com/watch?v=GoqD5kQs8nM&t=1s). Imitated R2-D2 sounds were identified by comparing the spectrograms of the bird imitations to the spectrogram of multiple sequences of the model R2-D2 sounds. R2-D2 sound imitations were cut out of each recording and saved as individual wav files for further analyses. We selected the four most frequently imitated monophonic and multiphonic R2-D2 sound units for further analyses. Sounds were determined as either monophonic or multiphonic sounds by visually inspecting the spectrograms and listening to the sounds.

We used Luscinia (version 2.22.12.01.01) to compare imitation accuracy between parrots and European starlings (29). Each imitated sound was annotated in Luscinia on an elemental (single sounds) and unit level (consisting of multiple elements). We used Luscinia’s dynamic time warping algorithm to compare imitated R2-D2 units to the original model R2-D2 units. The following spectral temporal weightings were used (Table S1): time (5), peak frequency (1), fundamental frequency (1), peak frequency change (0.25), fundamental frequency change (0.25), gap between elements (0.5), PF norm (1), FF norm (1). We used an additional parameter (frequency bandwidth = 0.25) for the multiphonic unit “C” due to the complexity of this sound unit. This had no effect on the overall results of the study i.e. similar scores but more variation between individuals. The dynamic time warping algorithm tries to optimally align two time series based on five alignment points. Dissimilarity scores are then calculated based on the weightings of the features for each alignment. For more details see https://github.com/rflachlan/Luscinia/wiki/Time-Warping-Analysis. The lower the dissimilarity scores are the more accurate the imitation was. Dissimilarity scores calculated by Luscinia have been used in numerous studies to compare similarities between avian songs and calls (35-38).

### Statistical Analysis

We created a Linear Mixed Model to test differences in multiphonic and monophonic R2-D2 unit imitation accuracy using the “lme4” package version 1.1-34 in R (version 2023.06.1) (40). Dissimilarity score was normalized to a mean of 0 and standard deviation of 1 and set as response variable. Animal ID was set as random effect and sound type (factor: multiphonic or monophonic) and starling (factor: starling or no-starling) were set as fixed effect together with an interaction term between the latter two. A second model was created that was similar to the first one but contained species as a fixed effect instead of starling to test if there were any differences between species in imitation accuracy. We assumed the data was normally distributed based on visual inspection of the residuals and high sample size (n = 101). An ANOVA test confirmed that the full model had a better fit than the null model (ANOVA: *χ*^2^ = 148.038, *df* = 2, *p* < 0.001). Finally, a Post-hoc Tuckey test was conducted to make comparisons between the factor levels of the interaction between the factors sound type and starling using the “emmeans” package version 1.8.9 (41). All statistical analyses were conducted in R.

### Dendrograms

We created dendrograms to compare species imitation of R2-D2 units to each other for both monophonic and multiphonic sound units separately. First, average dissimilarity scores were calculated between each species and the R2-D2 model sounds. Averages were calculated based on all four monophonic and all four multiphonic sound units. Next, we transformed the average dissimilarity scores into an Euclidean score matrix using the “dist” function in R. Lastly, we created dendrograms based on either all monophonic or multiphonic units using the “ggdendro” package version 0.2.0 in R (42).

## Acknowledgments

We thank Robert Lachlan for his help with using the dynamic time warping algorithm in Luscinia. We thank Carel ten Cate for the interesting discussions and suggestions for the analysis. This research was funded by FWF YIRG grant 10.55776/ZK66 & NWO XS grant 406.XS.03.074 (both awarded to MS).

