## Supplemental Information for "Artoo-Detoo: What imitating a Star Wars droid reveals on allospecific vocal imitation in parrots and starlings"

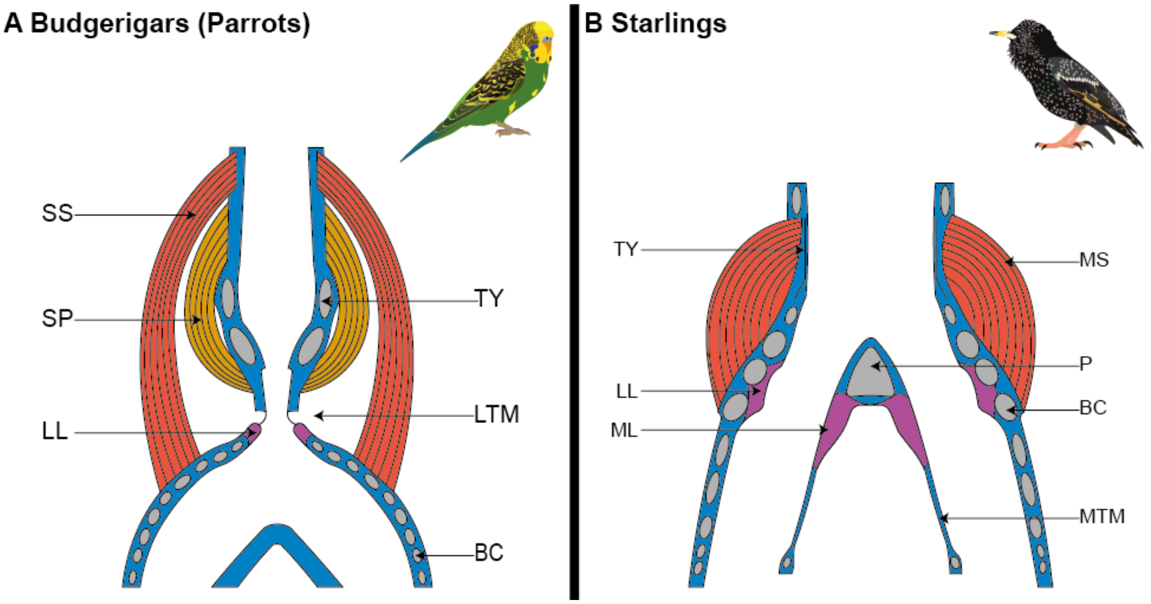


Fig. S1. **Anatomical differences between syrinxes of parrots and starlings**
(*A*) Schematic overview of the syrinx of a budgerigar as a representative of parrots. Abbreviations: SS, m. syringealis superficialis; SP, m. syringealis profundus; LL, lateral labia; TY, tympanum; LTM, lateral tympaniform membrane; BC, bronchial cartilage. Based on Abdel-Maksoud et al. (1) and partly modified from Larsen & Goller. (2).
(*B*) Schematic overview of a starlings' syrinx. Abbreviations: TY, tympanum; LL, lateral labia; ML, medial labia; MS, intrinsic syringeal muscles; P, pessulus; BC, bronchial cartilage; MTM, median tympaniform membrane. Note the lateral (LL) and medial labia (ML) that are part of the syrinx of the starling and play an important role in biphonation, are different in the budgerigar's syrinx without the presence of the medial labia. Modified from Prince et al. (3).

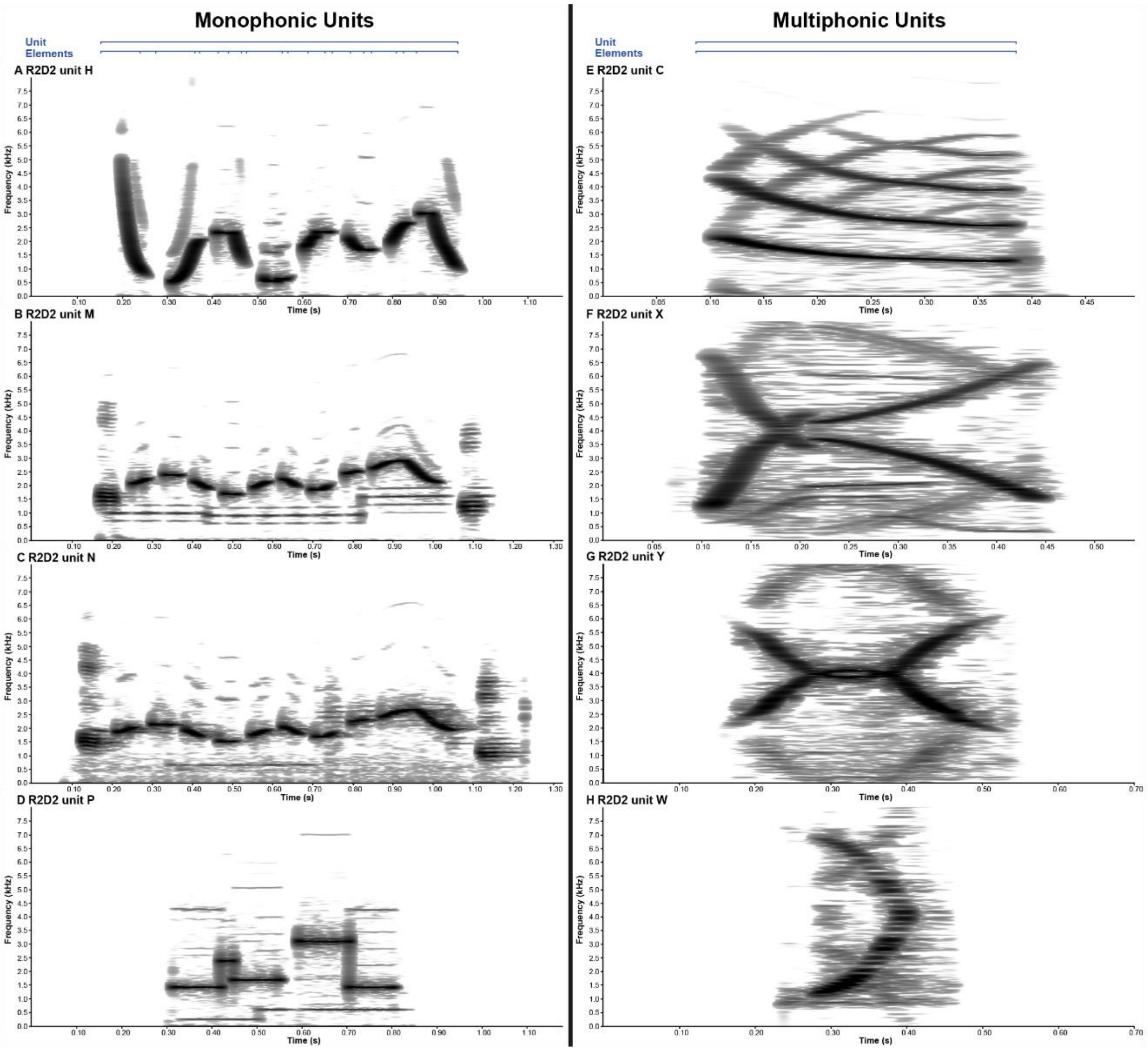


Fig. S2. **Spectrograms of R2-D2 model sounds**

R2-D2 sounds consist of elements (single sounds) and units (combination of one or more elements). Spectrograms of monophonic R2-D2 units are shown on the left side of the figure. (*A*) Unit “H”. (*B*) Unit “M”. (*C*) Unit “N”. (*D*) Unit “P”. Note that unit “M” and “N” look similar, but are different in the number of frequency modulations (i.e. maximum fundamental frequencies are 3.0 and 2.7 kHz for unit “M” and “N”, respectively). On the right side the multiphonic units are shown: (*E*) Unit “C”. (*F*) Unit “X”. (*G*) Unit “Y”. (*H*) Unit “W” (see Movie S1 for sound examples).

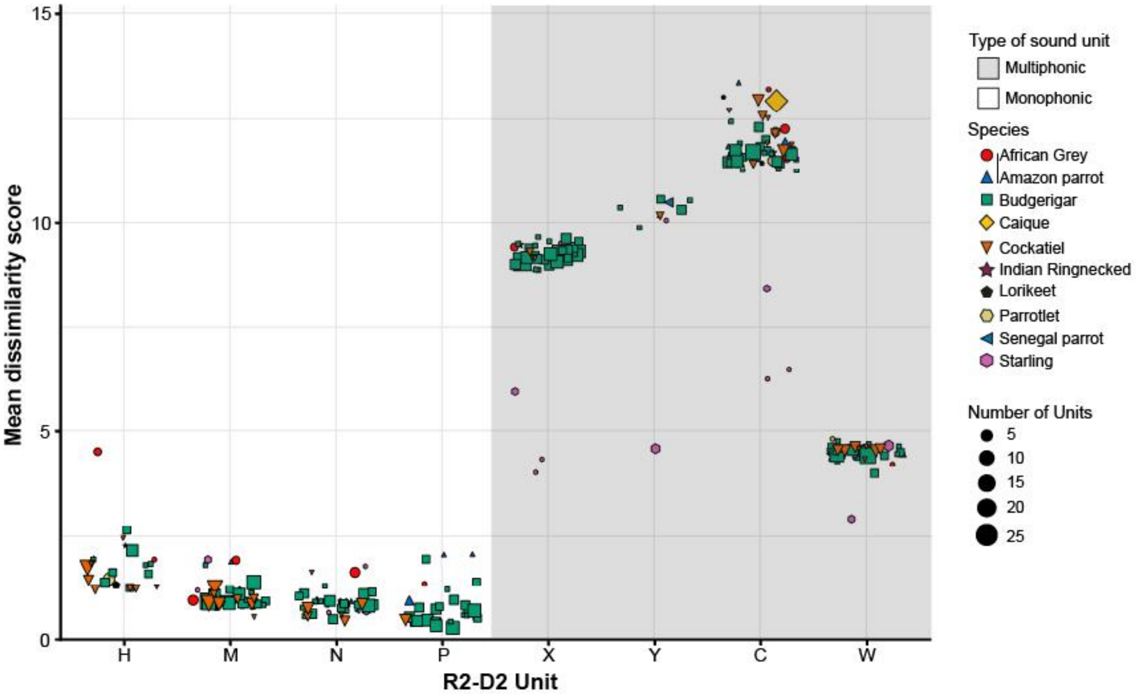


Fig. S3. **Imitation accuracy of R2-D2 units varied between individuals within species**
Mean dissimilarity scores per unit for each individual are shown. Dissimilarity scores varied between each unit but also within each species. Colors and shapes indicate the different species. Scores in the white and grey background correspond to monophonic and multiphonic units, respectively. Sizes of dots are related to the number of imitated units per individual.

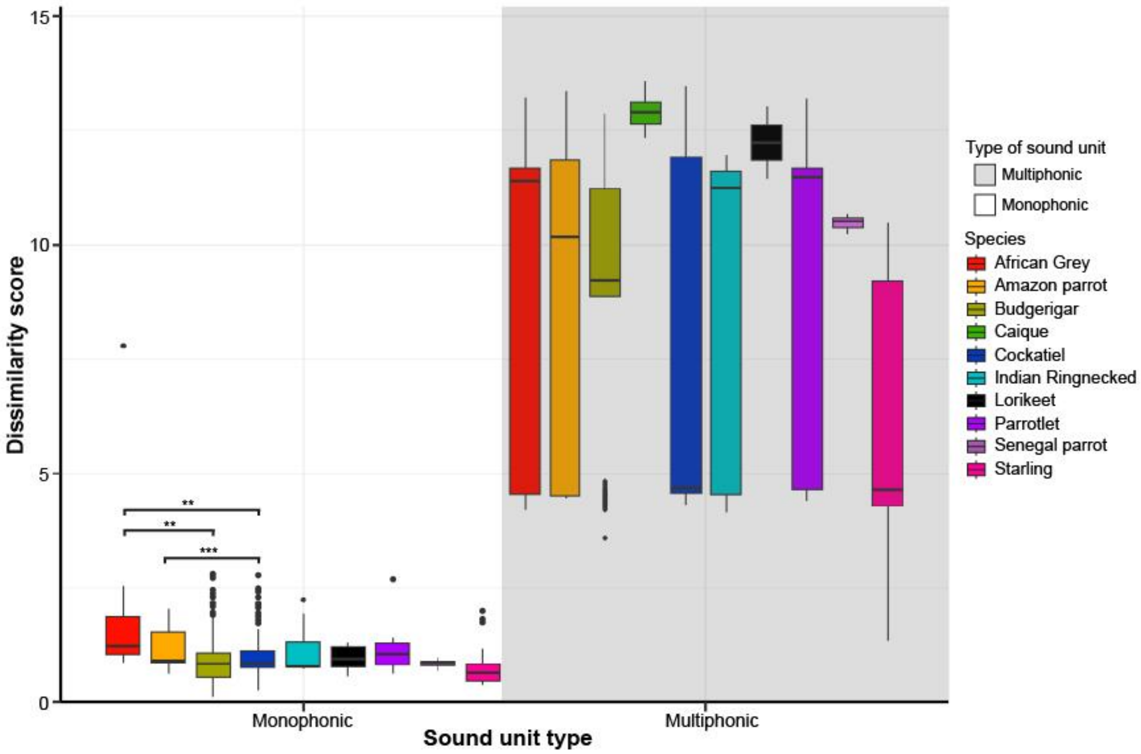


Fig. S4. **Imitation accuracy varied between species for both types of units**
Parrots imitate multiphonic units less accurately than starlings do. Parrots and starlings vary in how accurately they imitate monophonic units. Medians, inter-quartile ranges as well as minima and maxima are represented by horizontal lines, boxes and whiskers, respectively. Dissimilarity scores were combined for all monophonic and all multiphonic units separately. Colors correspond to dissimilarity scores for each species. Stars indicate significance levels (** $P < 0.01$, *** $P < 0.001$).

**Table S1. Spectral-temporal weightings for dynamic time warping in Luscinia**

| Spectral-temporal parameter | Weighting |
|---|---|
| Time | 5 |
| Peak frequency | 1 |
| Fundamental frequency | 1 |
| Peak frequency change | 0.25 |
| Fundamental frequency change | 0.25 |
| Gap between elements | 0.5 |
| Peak frequency normalized | 1 |
| Fundamental frequency normalized | 1 |
| Frequency bandwidth[a] | 0.25 |

[a] Only used for unit “C” due to complexity of the unit

**Table S2. Results Linear Mixed Model**

| Fixed effect | Estimate | SE | Min. estimate | Max. estimate | T-value | P-value |
|---|---|---|---|---|---|---|
| Intercept | 0.057 | 0.080 | -0.1000 | 0.214 | 0.708 | 0.481 |
| Starling: Starling | 0.469 | 0.327 | -0.170 | 1.108 | 1.433 | 0.154 |
| Type: Multiphonic | 0.141 | 0.066 | 0.012 | 0.269 | 2.140 | **0.033** |
| Starling*Multiphonic | -3.240 | 0.293 | -3.815 | -2.648 | -11.042 | **< 0.001** |

**Table S3. Result of Post-Hoc test**

| Fixed effect | Estimate | SE | Min. estimate | Max. estimate | T-value | P-value |
|---|---|---|---|---|---|---|
| No-starling Monophonic – Starling Monophonic | -0.469 | 0.328 | -1.318 | 0.380 | -1.431 | 0.482 |
| No-starling Monophonic – No-starling Multiphonic | -0.141 | 0.066 | -0.310 | 0.029 | -2.136 | 0.143 |
| No-starling Monophonic – Starling Multiphonic | 2.630 | 0.284 | 1.892 | 3.369 | 9.271 | **< 0.001** |
| Starling Monophonic – No-starling Multiphonic | 0.328 | 0.328 | -0.521 | 1.178 | 1.001 | 0.749 |
| Starling Monophonic – Starling Multiphonic | 3.099 | 0.286 | 2.362 | 3.837 | 10.820 | **< 0.001** |
| No-starling Multiphonic – Starling Multiphonic | 2.771 | 0.284 | 2.032 | 3.510 | 9.759 | **< 0.001** |

**Movie S1 (Model_Spectrograms_Final.MP4).** Rolling spectrogram of the four monophonic R2-D2 units "H", "M", "N" and "P" followed by the four multiphonic units "C", "X", "Y", "W". Created with the Dynaspec R package (4).

**Movie S2 (Imitation_Spectrograms_Final.MP4).** Rolling spectrogram of first the monophonic R2-D2 unit "N" followed by imitations of a starling and budgerigar, respectively. After the monophonic sounds an example of the multiphonic R2-D2 unit "X" is played back followed by imitations of a starling and budgerigar, respectively. Created with the Dynaspec R package (4).
